# A sex hormone-BDNF-TrkB axis directs sympathetic innervation in the mouse mammary gland

**DOI:** 10.1101/2025.10.18.683256

**Authors:** Subhajit Maity, Akshita Krishnan, Kloma T. Cardoza, Sami M. Frascoli, Darshan M. Joshi, Achira B. Shah, Dibyo Maiti, Purna A. Joshi

## Abstract

Stromal-epithelial interactions underlie fundamental tissue morphogenesis in normal development, tissue regeneration and cancer. Peripheral nerves are increasingly implicated as important stromal drivers of normal and malignant epithelial tissue biology. In the mammary gland, epithelial cell fate is highly dependent on stromal cues during distinct stages of postnatal growth which are triggered by ovarian sex hormones estrogen and progesterone. However, the effects of sex hormones on peripheral nerves in the postnatal mammary gland is largely unknown. Here, we uncover extensive changes in peripheral sympathetic innervation in the mammary stromal microenvironment during puberty and pregnancy which are periods of active postnatal mammary epithelial growth. We find that sex hormones induce sympathetic axonal branching through an intricate hormone-epithelial-nerve cross-talk. Specifically, estrogen and progesterone stimulate the expression of brain-derived neurotrophic factor (BDNF) in mammary hormone receptor-expressing luminal epithelial cells which acts on Tropomyosin receptor kinase B (TrkB)- expressing sympathetic nerves to activate BDNF-TrkB signaling. Our findings illustrate a previously unrecognized capacity of hormone-sensing epithelial cells to modulate sympathetic innervation, providing a framework for understanding nerve dynamics during tissue regeneration and cancer.

## Introduction

The stromal compartment of epithelial tissues such as the mammary gland is composed of diverse cellular elements, including fibroblasts, immune cells, endothelial cells, adipocytes, and peripheral nerves[1, 2]. Although peripheral nerves are observed to innervate the epithelial microenvironment, systemic and local signals that control peripheral innervation in postnatal tissues are not well defined. High peripheral nerve density, especially relating to sympathetic nerves, is associated with poor patient prognosis in cancers including breast cancer [3, 4].

Deciphering how peripheral innervation is dictated in the normal mammary gland will provide insight into their aberrant regulation in breast cancer.

The mammary gland is a vital organ that supports reproduction by producing milk to nourish the young. Unlike most other organs, the mammary gland undergoes massive postnatal tissue remodeling during critical stages including puberty, pregnancy, lactation, involution and estrous cycles in mouse or menstrual cycles in human[5]. These changes involve intricate epithelial reorganization driven by tissue-resident stem and progenitor cells in concert with intrinsic factors as well as extrinsic signals derived from the niche[6, 7]. Sex hormones, particularly estrogen and progesterone, are pivotal regulators of stem/progenitor cells and consequently postnatal mammary epithelial morphogenesis[6, 8–10]. While peripheral nerves are known to innervate the mammary stroma, their dynamics during postnatal mammary epithelial growth have remained elusive.

In this study, we show that sympathetic nerve density increases in the mammary gland during pubertal development and pregnancy. We observe that estrogen and progesterone stimulation increases brain-derived neurotrophic factor (BDNF) in mammary luminal epithelial cells which in turn recruits sympathetic nerves by activating BDNF-TrkB signaling.

## Results

### Peripheral and sympathetic innervation exhibits dynamic changes during postnatal mammary morphogenesis

First, we set out to determine whether peripheral nerve density undergoes dynamic changes during postnatal mammary epithelial remodeling. We quantified nerve fiber density in wild-type mice at four distinct developmental stages: prepubescent (two weeks), puberty onset (three weeks), mid-puberty (six weeks), and adult (twelve weeks) **(Figure 1A, 1B)**. Immunostaining and analysis for Protein Gene Product 9.5 positive (PGP9.5^+^) nerve fibers in the mammary gland indicated that total peripheral innervation undergoes dynamic changes during pubertal development. We observed that PGP9.5^+^ nerve fiber density is highest at the onset of puberty **(Figure 1C, 1F)** when the mammary gland is highly proliferative **(Figure 1D)**. Further analysis for Tyrosine Hydroxylase positive (TH^+^) nerve fibers marking sympathetic nerves indicates that sympathetic nerve density peaks similarly at puberty onset compared to the prepubescent stage **(Figure 1E, 1G)**. Interestingly, sympathetic nerve density in the mammary gland significantly drops at mid-puberty and remains reduced in adult glands.

**Figure 1:**
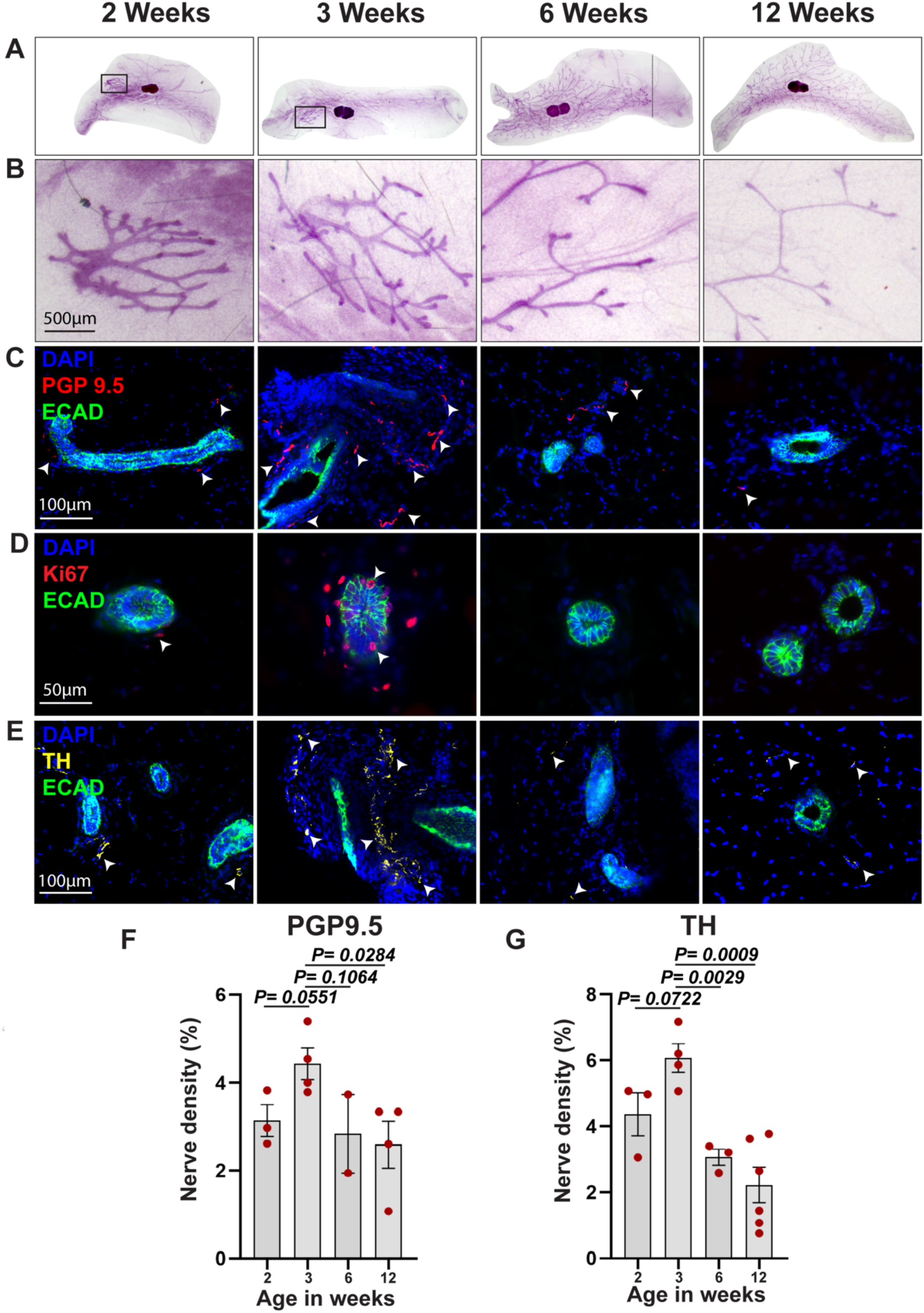
Peripheral and sympathetic nerve innervation undergo dynamic changes during postnatal mammary branching morphogenesis. (A,B) Carmine alum-stained mammary gland whole mounts showing epithelial branching distribution during postnatal development. (**C)** Immunofluorescent staining of mammary gland tissue sections with total peripheral nerve marker Protein gene product 9.5 (PGP9.5), (**D)** Sympathetic nerve marker Tyrosine hydroxylase (TH), and epithelial marker E-cadherin (ECAD). Arrowheads indicate nerve fibers. (**E)** Immunofluorescent staining with Ki-67 to visualize proliferating cells. (**F)** Peripheral nerve density, and **(G)** sympathetic nerve density in the mammary gland during development. DAPI stains nuclei. Each dot represents one biological replicate. Three to five whole-side images were analyzed for each biological replicate and their mean was plotted for each sample. Data represent the mean ± s.e.m.

Next, we probed innervation changes during pregnancy given that it is characterized by a profound expansion in mammary epithelia, culminating in the genesis of milk-secreting lobuloalveolar structures. We subjected adult wild-type female mice to timed pregnancy at 10 weeks of age following recommendations from the Jackson Laboratory. Mice were sacrificed mid-pregnancy (day 12.5) and mammary glands were harvested for subsequent comparative analysis with those from nulliparous female mice at the resting estrus stage of the cycle. As expected, mid-pregnancy mammary glands exhibited marked lobuloalveologenesis (**Figure 2A**). Immunofluorescence staining and quantification showed that PGP9.5^+^ **(Figure 2B, C)** and TH^+^ **(Figure 2D, E)** nerve fiber density is significantly higher in the mammary glands of mid-pregnant females as compared to nulliparous estrus controls. Taken together, these data show that peripheral and sympathetic nerve density demonstrate variable changes in the mammary gland during puberty and mid-pregnancy. Intriguingly, this increase in nerve density coincides with high levels of sex hormone-mediated epithelial branching morphogenesis in the postnatal mammary gland.

**Figure 2:**
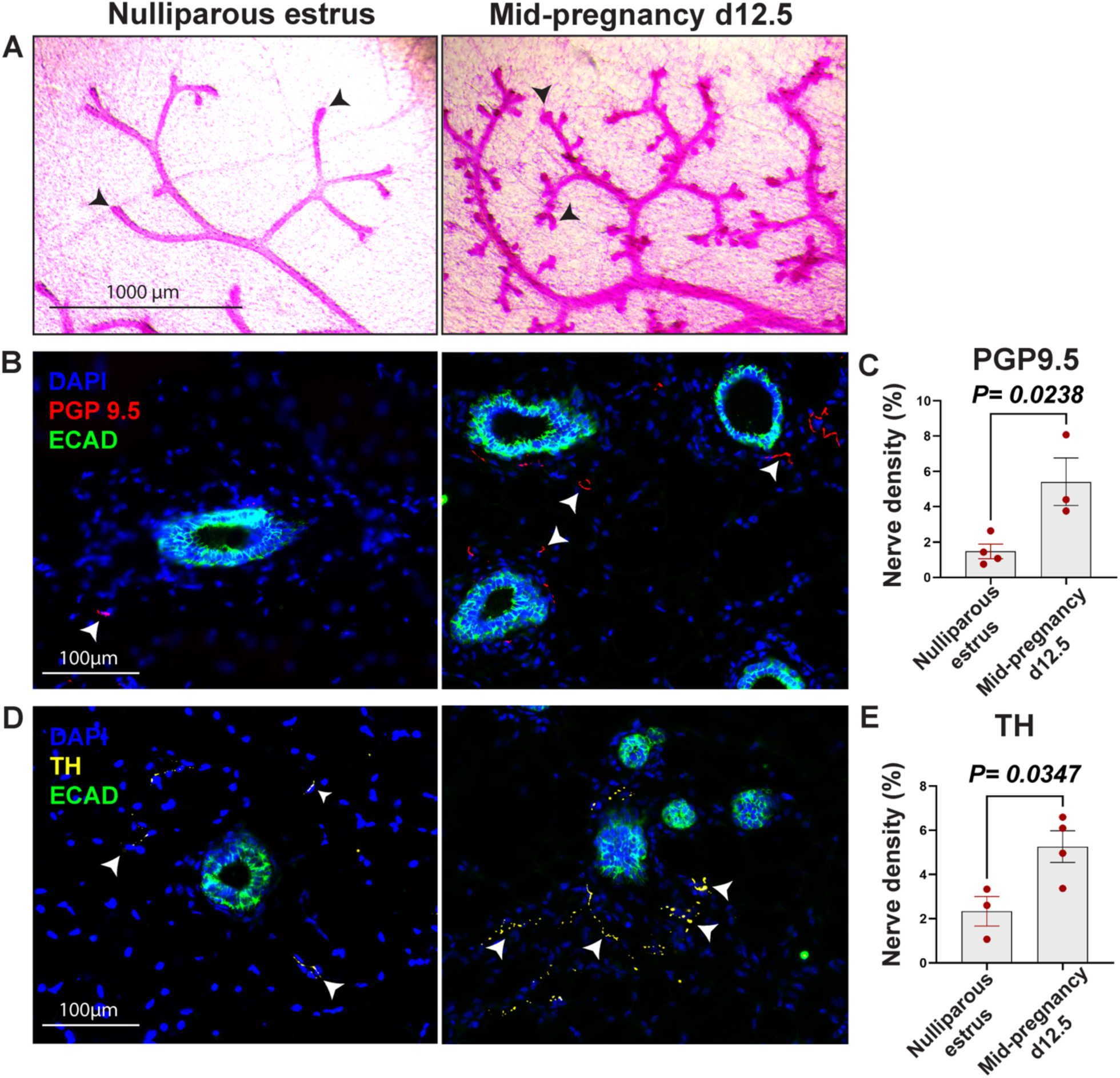
Enhanced peripheral and sympathetic nerve innervation in the mammary gland during lobuloalveolar differentiation in pregnancy. **(A)** Carmine alum-stained mammary gland whole mounts showing epithelial branching in nulliparous estrus phase and pregnancy. Arrowheads indicate ductal termini in resting nulliparous glands and proliferative alveolar structures at mid-pregnancy d12.5. Immunofluorescence staining on mammary gland sections with **(B, C)** peripheral nerve marker PGP9.5 and epithelial marker ECAD, **(D, E)** sympathetic nerve marker TH and epithelial marker ECAD. Arrowheads indicate nerve fibers. Quantified nerve density is plotted where each dot represents one biological replicate. For each replicate, 3-5 whole-slide images were quantified and their mean was plotted. Data represent the mean ± s.e.m.

### Sex hormones drive sympathetic nerve innervation in the mammary gland

Given the observed increase in both peripheral and sympathetic nerve fiber density in the mammary gland at the onset of puberty when estrogen levels are high, and during pregnancy when both estrogen and progesterone levels rise, we sought to investigate whether sex hormones could regulate changes in peripheral nerve innervation. To test this hypothesis, we treated adult 10-week-old wild-type female mice with 17β-estradiol and progesterone (EP) to induce mammary epithelial regeneration and alveologenesis as previously described[8]. As anticipated, mammary glands isolated from EP-treated mice had extensive branching and alveolar expansion **(Figure 3A)**. Notably, we found that PGP9.5^+^ peripheral **(Figure 3B, 3C)** and TH^+^ sympathetic nerve fiber density **(Figure 3D, 3E)** were significantly elevated in the stroma of EP-treated mice compared to untreated controls. These observations suggest that sex hormones augment sympathetic innervation in the mouse mammary gland during epithelial tissue morphogenesis.

**Figure 3:**
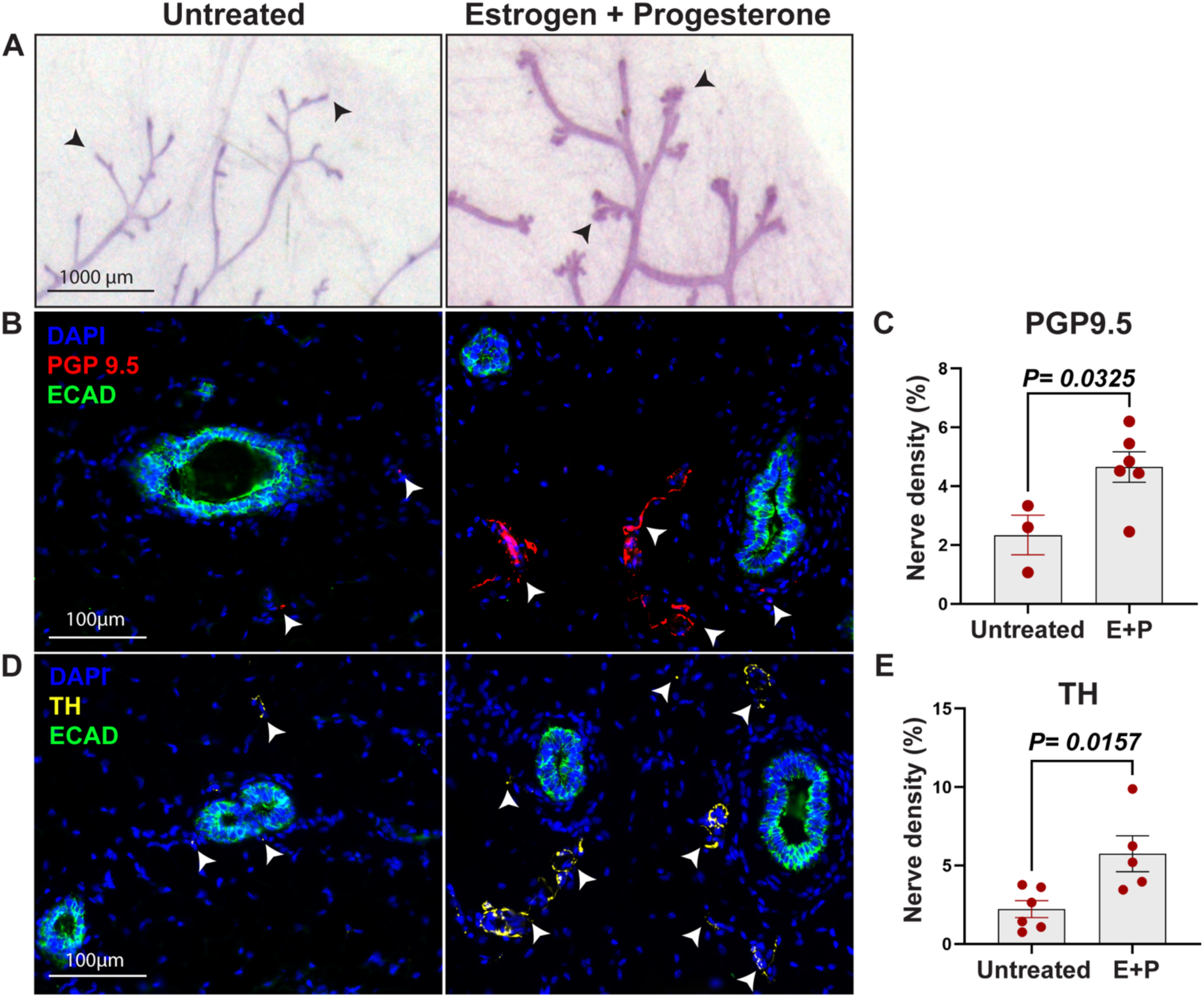
Estrogen and Progesterone treatment increases peripheral and sympathetic nerve density in the mammary microenvironment. **(A)** Carmine alum-stained mammary gland whole mounts showing epithelial branching in untreated and estrogen + progesterone (E+P)- treated samples. Arrowheads indicate ductal termini in untreated mice and alveolar structures in E+P-treated mice. Immunofluorescence staining on mammary gland sections with **(B, C)** peripheral nerve marker PGP9.5 and epithelial marker ECAD, **(D, E)** sympathetic nerve marker TH and epithelial marker ECAD. Arrowheads indicate nerve fibers. Quantified nerve density is plotted where each dot represents one biological replicate. For each replicate, 3-5 whole-slide images were quantified, and their mean was plotted. Data represent the mean ± s.e.m.

### BDNF is differentially upregulated in hormone receptor-enriched mammary epithelial cells during glandular expansion

Based on our finding that mammary nerve density is heightened following sex hormone exposure, we interrogated whether this increase in nerve density is due to axonal restructuring mediated by neurotrophic factors. First, we analyzed our previously published bulk RNA sequencing dataset (GSE123714) in which mammary cells were sorted from untreated or EP-treated mice[11]. We specifically analyzed mammary luminal epithelial cells, and segregated them based on their expression of *Esr* (estrogen receptor) and *Pgr* (progesterone receptor) to denote luminal populations expressing low (HR^lo^) and high (HR^hi^) levels of sex hormone receptors **(Figure 4A, B)**. As expected, *Ly6a* (Sca1)[12] and *Tnfsf11* (RANKL)[8] was higher in the HR^hi^ luminal subset compared to the HR^lo^ subset following EP stimulation **(Figure 4C, D)**. We then examined the expression of neurotrophic factors in these luminal subsets and detected a striking increase in the expression of brain-derived neurotrophic factor (*Bdnf*) in the HR^hi^ mammary luminal epithelial subset of EP-treated mice compared to untreated controls **(Figure 4E)**. To further validate this increase in luminal *Bdnf* in response to sex hormones, we sorted mammary luminal epithelial cells from untreated and EP-treated wild-type female mice **(Supplementary Figure 1)** and quantified their *Bdnf* transcript levels. Similar to our observations using the RNA-seq data, *Bdnf* expression was higher in EP-treated luminal cells compared to untreated controls **(Figure 4F)**. We then analyzed mammary tissue sections from these mice for BDNF protein by immunofluorescence staining, and found that mean fluorescence intensity levels of BDNF were significantly higher in hormone-treated samples as compared to untreated controls **(Figure 4G, H)**. Thus, our data confirm enhanced BDNF induction in the mammary luminal epithelial compartment, predominately in the hormone-sensing subset, in response to sex hormones during epithelial regeneration.

**Figure 4:**
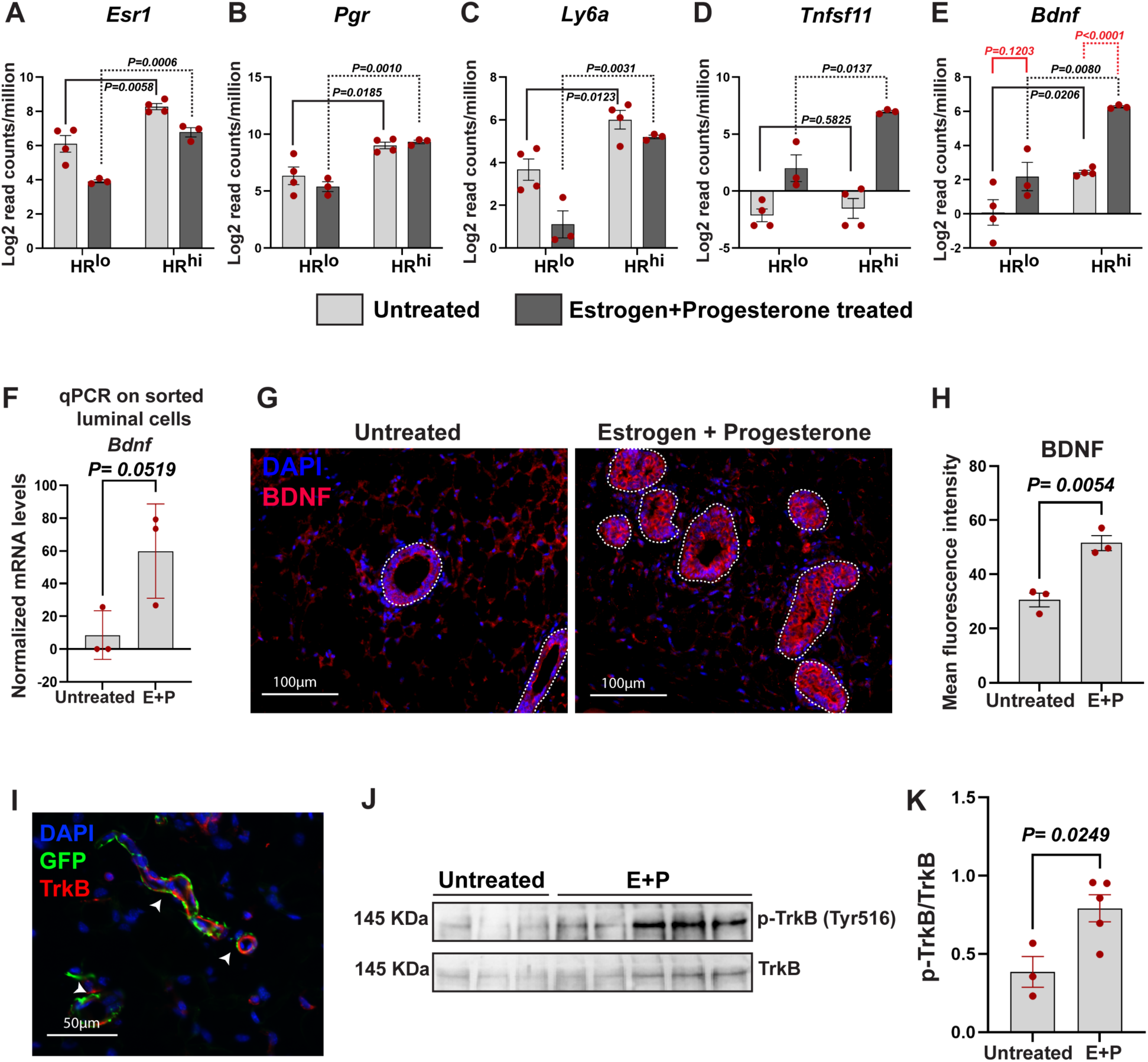
Sex hormones induce a neurotrophic program in the mammary gland via luminal epithelial cells. (A-E) Analysis of bulk RNA-seq data[11] showing increased *Bdnf* expression in luminal mammary epithelial cells that are enriched in hormone receptors. **(F)** qPCR analysis of sorted luminal mammary epithelial cells from EP-treated mice showing increased *Bdnf* expression compared to untreated controls. **(G, H)** Immunofluorescent staining of mammary gland sections from untreated or EP-treated wild-type mice for BDNF and DAPI. For each replicate, quantification was performed by analyzing 5 fields from each of 5 sections per replicate (25 images in total per replicate) and their average was plotted as one data point. Data represent the mean ± s.e.m. **(I)** Mammary gland sections from TH-2A-CreERT2 Rosa^mT/mG^ mice probed for GFP, TrkB, and DAPI, showing GFP⁺TrkB⁺ nerve fibers (arrowheads), indicating expression of TrkB receptors on TH^+^ sympathetic nerves. **(J, K)** Western blot of mammary gland tissues from untreated or EP-treated wild-type mice probed for phosphorylated TrkB (pTrkB) and total TrkB. Data represent n=3 for untreated and n=5 for E+P-treated groups with mean ± s.e.m.

### Estrogen and progesterone activate BDNF-TrkB signaling in the mammary gland

BDNF is a well-known chemotactic regulator of sympathetic neurons expressing the cognate receptor TrkB[13]. In mammary tissue sections from adult TH-2A-Cre^ERT2^ Rosa^mT/mG^ reporter mice, we observed the localization of TrkB to GFP^+^ nerve fibers representing TH+ sympathetic neurons **(Figure 4I)**, affirming the presence of the BDNF receptor on sympathetic nerves innervating the mammary gland. Next, we wanted to assess whether a sex hormone-driven increase in luminal BDNF expression leads to activation of BDNF-TrkB signaling. To investigate this, we performed immunoblotting for TrkB and 6hosphor-TrkB (p-TrkB) using mammary tissue protein harvested from untreated and EP-treated mice. We observed a marked increase in TrkB phosphorylation levels as demonstrated by an increase in p-TrkB/TrkB ratio in mammary glands of hormone-treated mice compared to untreated controls **(Figure 4J, K)**. Collectively, these data show that estrogen and progesterone can activate BDNF-TrkB-mediated neurotrophic signaling in the mammary gland during hormone-driven tissue growth.

## Discussion

Unlike the central nervous system, the dynamics of nerve innervation in peripheral tissues have been historically understudied. While previous studies identified a sexually dimorphic pattern of innervation in the embryonic mammary gland[2, 14], innervation patterns and underlying mechanisms during postnatal mammary tissue morphogenesis are not understood. In this study, our findings reveal that peripheral and sympathetic nerve innervation is highly adaptable and responsive to the hormonal milieu in the postnatal mouse mammary gland. The marked increase in sympathetic nerve density during critical proliferative windows of pubertal development and pregnancy-induced regeneration of milk-secreting alveolar structures, as well as following sex hormone stimulation, underscores a fundamental sex hormone-triggered sympathetic nerve patterning program. We identify BDNF as a candidate mediator of this program wherein sex hormones induce the expression of this neurotrophic factor primarily in hormone-sensing mammary luminal epithelial cells. We note that sympathetic nerves in the mammary gland express TrkB receptors for BDNF and that sex hormone stimulation does indeed lead to TrkB receptor phosphorylation and activation of the BDNF-TrkB signaling pathway in the mammary gland **(Figure 5)**.

**Figure 5:**
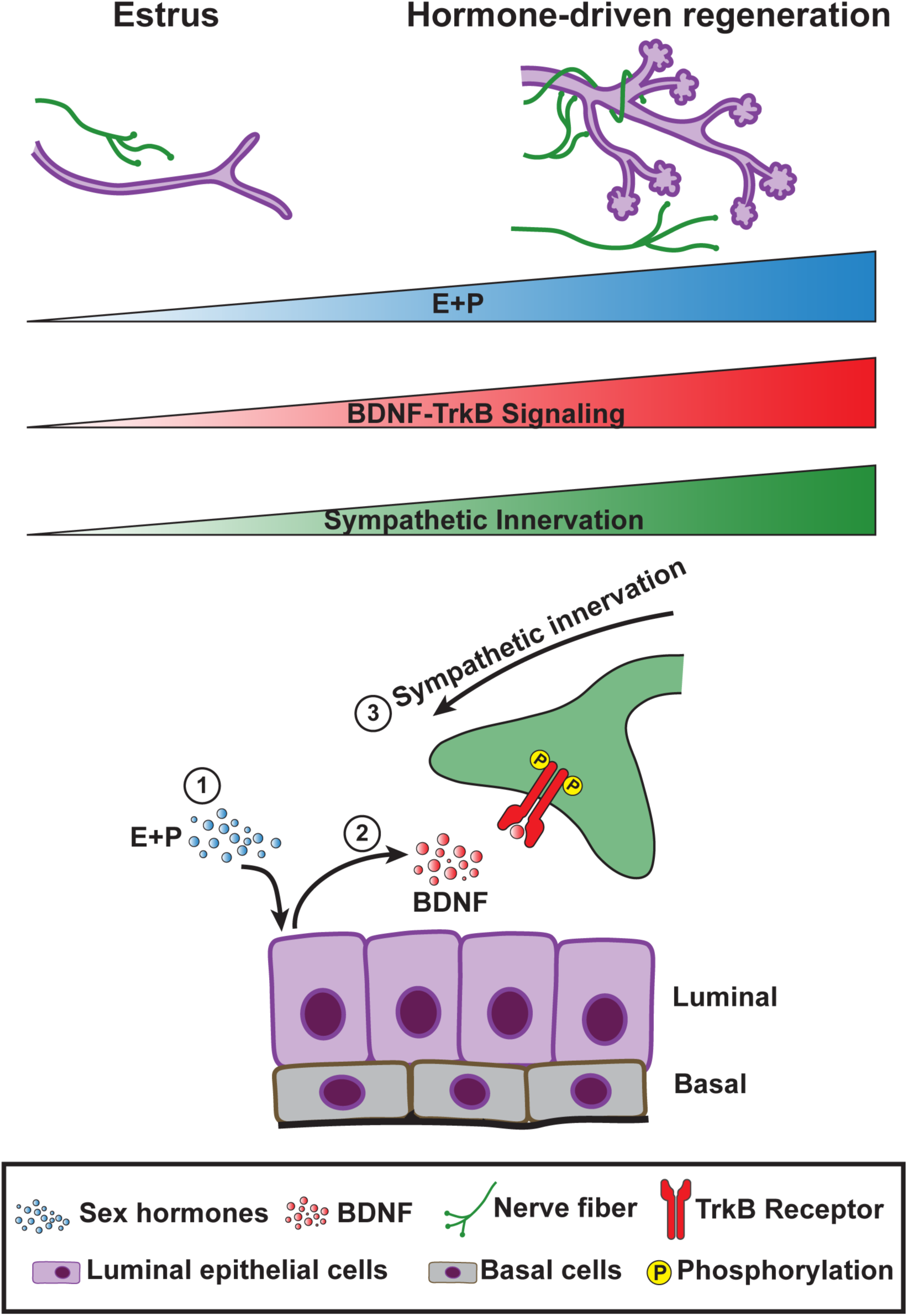
Model of sex hormone-mediated neurotrophic signaling during mammary epithelial regeneration. Sympathetic nerve innervation in the mammary gland is highly responsive to sex hormones estrogen and progesterone (EP) during lobuloalveologenesis in the mammary gland. Specifically, hormone-triggered BDNF secretion from luminal epithelial cells leads to activation of BDNF-TrkB signaling, culminating in increased sympathetic innervation.

Sex hormones are well known to drive breast cancer growth and progression[15–17]. High sympathetic nerve density in mammary tissues is associated with poor prognosis in breast cancer patients[3], and increased sympathetic activity has been shown to enhance metastatic potential of breast cancer cells[4]. Although sex hormones and sympathetic nerve density are independently linked to breast cancer progression, how sex hormones may influence sympathetic nerve dynamics in mammary tissue was not known. Our work infer a capacity of sex hormones to induce a neurotrophic program driving sympathetic innervation which provides insight into how sex hormones may control nerve density in pathological tissue states such as cancer. While much has been learned from studies delving into embryonic mammary innervation[2, 14], there is a paucity of knowledge about innervation during key postnatal hormonally-controlled developmental stages like pregnancy and puberty which are crucial for understanding growth and patterning processes involved in not just normal biology but also disease. Neurotrophic factors like BDNF have been reported to be present at high levels during specific phases in the ovine mammary gland[18], and its systemic levels are linked to mouse mammary epithelial development[19], but the effects of such factors on mammary sympathetic innervation remained elusive until now. Our observations fill this important gap by identifying a unique sex hormone-driven epithelial-nerve crosstalk in which epithelial cells serve as an intermediary that helps in recruiting sympathetic nerves via secreting neurotrophic factor BDNF, where sympathetic nerves likely contribute to epithelial growth and remodeling.

The mammary microenvironment contains diverse classes of neurotrophic cues such as Netrin, semaphorins and others, in addition to multiple types of peripheral nerves including sympathetic and sensory nerves, each of which may contribute uniquely to epithelial remodeling and subsequently influence cell fate and cancer outcome. By defining a sex hormone-BDNF-TrkB mechanism for postnatal peripheral sympathetic innervation, our work establishes a framework that can be applied to systematically explore the broader network of neurotrophic factors and nerve subtypes in epithelial tissues beyond the mammary gland that may work in concert with epithelial cells to likely influence tissue biology. Such efforts may uncover a complex circuitry of cellular candidates that can be targeted to improve the outcome for diseases such as cancer.

## Methods

### Mice

C57BL/6J (strain: 000664), TH-2A-Cre^ERT2^ (strain: 025614), ROSA^mTmG^ (strain: 007676) mice were procured from Jackson Laboratories and housed on a 12-hour light and 12-hour dark cycle with food and water ad libitum. Untreated female mice were screened to confirm their estrus stage by vaginal cytology. All procedures were approved by the Institutional Animal Care and Use Committee at The University of Texas at Dallas.

### Timed pregnancy

10-week-old adult wild-type female mice were subjected to timed pregnancy according to recommendations from The Jackson Laboratory. Females with synchronized estrus cycles were mated with a male mouse and inspected for the presence of a vaginal mucus plug the following morning to confirm mating which was then designated as day 0.5 of pregnancy. Mice were regularly monitored for weight gain to confirm pregnancy. Pregnant females were sacrificed at day 12.5 of pregnancy, and mammary glands were harvested for analysis.

### Hormone treatment

Adult 10-week-old wild-type C57BL/6J female mice were anesthetized using isoflurane and an incision was made between the ears and the shoulder on the dorsal side of the mouse, and a 14- day slow-release pellet containing 17ß-Estradiol (0.14mg) + Progesterone (14mg) was implanted subcutaneously. Subsequently, the incision was closed using a surgical clip, and mice were analyzed after 14 days of hormone treatment.

### Tissue processing, immunofluorescent staining, imaging and analysis

Harvested mammary glands were fixed with 4% paraformaldehyde for 2 hours at room temperature, followed by three washes with 1X PBS (5 minutes each) on a nutating mixer. Tissues were then cryopreserved in 30% sucrose in 1X PBS at 4°C for one week on a nutating mixer. Sucrose was replaced with a fresh solution after the first 3 days. Subsequently, tissues were embedded in Optimal Cutting Temperature (OCT) medium and frozen at -80°C. Tissue sections (10μm) were made from OCT-embedded tissues using a Leica CM1950 cryostat and stored in - 80°C. Prior to staining, sections were air dried and hydrophobic barriers were drawn around sections using a barrier pen. Sections were then washed with 1X PBS and treated with permeabilization and blocking buffer containing 5% normal donkey serum, 1% BSA, and 0.2% Triton-X in 1X PBS for one hour at room temperature followed by three washes with 1X PBS. Sections were subsequently stained with appropriate primary antibody cocktails, prepared in a blocking buffer containing 5% normal donkey serum and 1% BSA in 1x PBS, and incubated overnight at 4°C. The following day, primary antibodies were removed and sections were washed three times with 1X PBS and incubated with appropriate secondary antibody cocktails prepared in blocking buffer for one hour at room temperature. Sections were washed again and mounted with ProLong Gold containing DAPI mounting media, and imaged using either Echo Revolve, Zeiss Axioimager A2 or Evident Scientific VS2000 Research Slide Scanner. Images of whole sections were analyzed using Fiji or cellSens to determine total nerve density. Region of Interest (ROI) were drawn to mark the boundaries of mammary tissues. Single-channel images were used to measure DAPI, PGP9.5, and TH-stained areas within ROIs. Nerve density of calculated using the following formula.

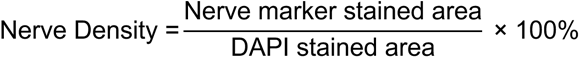

Alternatively, harvested fresh tissues were fixed in 4% PFA overnight at 4°C, followed by three washes in 1X PBS and storage in 70% ethanol at 4°C. Tissues were then processed using a 12- hour preset protocol in the Leica ASP300 S fully enclosed tissue processor and embedded in paraffin using the Leica ARCADIA C and H embedder. Next, 5-10μm sections were obtained using a Leica RM2235 microtome. Tissue sections were first deparaffinized in xylene for 20 minutes, followed by washes of 100% ethanol (2 washes) for three minutes each, 90% ethanol for three minutes, 70% ethanol for three minutes, and deionized water for three minutes. Next, deparaffinized and hydrated sample sections were subjected to heat-induced epitope retrieval using Tris-EDTA buffer (pH 9) at 110°C for 30 minutes. Sections were allowed to cool down for 30 minutes and washed with PBS for 5 minutes. Sections were permeabilized with 0.2% Triton-X in PBS for 5 minutes and rinsed twice with PBS. Next, to quench autofluorescence, samples were treated with 3% hydrogen peroxide for 15 minutes and rinsed twice with PBS. Sections were then blocked with 5% donkey serum, 100μg/ml saponin, 3% BSA in PBS for one hour, and rinsed with 1X PBS twice, following which appropriate primary antibodies were added and incubated overnight at 4°C. Primary antibodies were removed the next day and sections washed with 1x PBS, and secondary antibodies added for an hour at room temperature. Following removal of secondary antibodies and three 1x PBS washes for 5 minutes each, sections were mounted with Prolong Gold containing DAPI.

To quantify BDNF levels, ROIs were drawn around mammary epithelial ducts in BDNF-stained mammary gland sections, and the mean gray value was quantified as a measure of BDNF protein expression levels.

### Whole mount carmine alum staining

Harvested whole mammary glands were spread on a glass slide and fixed overnight at room temperature using Carnoy’s fixative (6 parts 100% ethanol, 3 parts chloroform, and 1 part glacial acetic acid). After, fixed samples were rehydrated in a gradually decreasing concentration of ethanol (70%, 50%, 30%, 10%) twice for 15 minutes each, followed by 5 minutes incubation in deionized water. Samples were subsequently incubated with carmine alum stain overnight at room temperature. The next day, samples were gradually dehydrated using an increasing concentration of ethanol (70%, 95%, 100%) twice for 15 minutes each. Tissues were left in xylene overnight to delipidate the tissue and mounted using Permount.

### Analysis of RNA-seq data

We have analyzed previously published bulk RNA sequencing data[11] with accession number GSE123714. Raw gene expression counts were normalized to counts per million (CPM) in R using the DGEList object to account for differences in sequencing depth across samples. CPM values were calculated by dividing raw counts by the total library size for each sample and multiplying by 1 million. Log2 transformation of the CPM values was then applied to stabilize variance and reduce skewness in the data. This normalization enabled the comparison of gene expression levels across samples while minimizing the influence of extreme count values.

### Tissue dissociation and fluorescence-activated cell sorting

Mammary glands were isolated from adult mice and digested for 2.5 hours at 37°C in DMEM-F12 media containing 750U/ml collagenase and 250 U/ml hyaluronidase. After 2.5 hours, dissociated tissues were vortexed and the cell suspension was washed with 1X Hanks Balanced Salt Solution (HBSS) containing 2% FBS (HF) to remove the digestion media. Next, samples were incubated with 0.8% ammonium chloride solution to lyse the red blood cells, further dissociated with 0.25% trypsin for 2 minutes, followed by 5U/ml dispase and 0.1mg/ml DNase I for 2 minutes. Digested samples were washed with HF and filtered through a 100mm strainer to acquire a single-cell suspension. For flow cytometry and sorting, cells were incubated with various antibodies to specific surface markers. A combination of PE-Cy7 conjugated anti-CD45, anti-Ter119 and anti- CD31 antibodies were used to label immune, erythrocyte and endothelial populations for exclusion by flow cytometry. The viability dye ZombieUV was used to mark live/dead cells. Mammary epithelial and stromal cell distributions were analyzed using APC-Cy7 conjugated anti-EpCAM and APC conjugated anti-CD49f antibodies . Luminal (ZombieUV^-^ CD45^-^ Ter119^-^ CD31^-^ EpCAM^hi^ CD49f^lo^), basal (ZombieUV^-^ CD45^-^ Ter119^-^ CD31^-^ EpCAM^lo^ CD49f^hi^) and stromal (ZombieUV^-^ CD45^-^ Ter119^-^ CD31^-^ EpCAM^-^ CD49f^-^) cells were sorted using a BD FACSAria^TM^ Fusion flow cytometer.

### RNA isolation and qPCR

Total RNA was isolated from sorted primary cells using Qiagen RNeasy Micro kit, and the cDNA first strand was synthesized from 10ng of total RNA using SMARTer^Ò^ PCR cDNA synthesis kit followed by cDNA amplification using the Advantage^Ò^2 PCR kit as per manufacturer’s protocol. Subsequently, qPCR was performed for genes of interest using Applied Biosystems QuantStudio 6 Flex and corresponding reagents following the manufacturer’s protocol. Data was analyzed to calculate and plot 2^-DDCt values.

### Protein isolation, SDS-PAGE and western blot analysis

Flash-frozen mammary glands were pulverized using Cellcrusher-mini. Lysis buffer containing 0.5% IGEPAL-CA630, protease and phosphatase inhibitor in TBS was added and the samples were vortexed on ice for 30 minutes in 5-minute intervals. Next, the lysates were ultrasonicated on ice using 10% amplitude 3 times for 5 seconds each. Subsequently, samples were centrifuged at 20,000g for 15 minutes at 4°C. Supernatants were collected and stored in -80°C. Proteins were quantified using the BCA assay kit following manufacturer’s protocol. Protein samples were mixed with Laemmli buffer and boiled at 98°C for 5 minutes and then centrifuged at maximum speed for 2 minutes. Following that, samples were resolved on a 10% polyacrylamide gel by SDS-PAGE and transferred to a PVDF membrane using the Bio-Rad Trans-Blot Turbo transfer system. After blocking with 5% BSA for 1 hour, appropriate primary antibodies are added and incubated overnight at 4°C. The next day, primary antibodies are washed off using three washes of TBST for 10 minutes each. Secondary antibodies were then added and incubated for 1 hour at room temperature and washed three times with TBST. Next, the blots were stained with MCE Ultra High Sensitivity ECL kit and chemiluminescence was visualized using the Invitrogen iBright 750 imaging system. Raw images were analyzed using Bio-Rad Image Lab to calculate the band intensity. Normalized band intensity was plotted and analyzed using GraphPad Prism.

### Statistical analysis

All statistical analyses were performed using GraphPad Prism and reported as mean ± standard error of the mean (SEM). Each n represents data acquired from one biological replicate. Comparison between two groups and statistical analysis was performed using Student’s t-test.

## Data availability

Raw data will be made available upon request.

## Materials

## Acknowledgements

This work is supported by UT Dallas Startup funds to PAJ. SM holds a Graduate Research and Cancer Education Fellowship from UT Dallas. The authors would like to sincerely thank Dr. Gregory Dussor, Dr. Theodore Price and Dr. Stephanie Shiers from the Department of Neuroscience at UT Dallas for their valuable input regarding neuronal marker selection. The authors are thankful to Dr. Jacob Henderson (Manager, UT Dallas FACS core) for his assistance with cell sorting and Dr. Joseph Lombardo (Manager, UT Dallas Imaging core) for his help with whole slide imaging and analysis.

## Author contributions

SM and PAJ conceptualized the study. Methodology was developed by SM and PAJ. SM, AK, SMF, KTC, and DMJ carried out the study. Data collection was performed by SM, AK, KTC, DMJ, and ABS, and formal analysis was conducted by SM, AK, KTC, and ABS. SM prepared the figures and DM contributed to generating the schematic illustration. The study was supervised and directed by PAJ. AK and ABS contributed to the writing of methodology. SM and PAJ wrote the manuscript.

## References

1. Wiseman BS, Werb Z. Stromal effects on mammary gland development and breast cancer. Science. 2002;296(5570):1046–9. doi: 10.1126/science.1067431. PubMed PMID: 12004111; PubMed Central PMCID: PMCPMC2788989.

2. Liu Y, Rutlin M, Huang S, Barrick CA, Wang F, Jones KR, et al. Sexually dimorphic BDNF signaling directs sensory innervation of the mammary gland. Science. 2012;338(6112):1357–60. doi: 10.1126/science.1228258. PubMed PMID: 23224557; PubMed Central PMCID: PMCPMC3826154.

3. Li D, Hu LN, Zheng SM, La T, Wei LY, Zhang XJ, et al. High nerve density in breast cancer is associated with poor patient outcome. FASEB Bioadv. 2022;4(6):391–401. Epub 20220303. doi: 10.1096/fba.2021-00147. PubMed PMID: 35664834; PubMed Central PMCID: PMCPMC9164247.

4. Sloan EK, Priceman SJ, Cox BF, Yu S, Pimentel MA, Tangkanangnukul V, et al. The sympathetic nervous system induces a metastatic switch in primary breast cancer. Cancer Res. 2010;70(18):7042–52. Epub 20100907. doi: 10.1158/0008-5472.CAN-10-0522. PubMed PMID: 20823155; PubMed Central PMCID: PMCPMC2940980.

5. Watson CJ, Khaled WT. Mammary development in the embryo and adult: new insights into the journey of morphogenesis and commitment. Development. 2020;147(22). Epub 20201115. doi: 10.1242/dev.169862. PubMed PMID: 33191272.

6. Joshi PA, Di Grappa MA, Khokha R. Active allies: hormones, stem cells and the niche in adult mammopoiesis. Trends Endocrinol Metab. 2012;23(6):299–309. Epub 20120519. doi: 10.1016/j.tem.2012.04.002. PubMed PMID: 22613704.

7. Fu NY, Nolan E, Lindeman GJ, Visvader JE. Stem Cells and the Differentiation Hierarchy in Mammary Gland Development. Physiol Rev. 2020;100(2):489–523. Epub 20190920. doi: 10.1152/physrev.00040.2018. PubMed PMID: 31539305.

8. Joshi PA, Jackson HW, Beristain AG, Di Grappa MA, Mote PA, Clarke CL, et al. Progesterone induces adult mammary stem cell expansion. Nature. 2010;465(7299):803–7. doi: 10.1038/nature09091. PubMed PMID: 20445538.

9. Asselin-Labat ML, Vaillant F, Sheridan JM, Pal B, Wu D, Simpson ER, et al. Control of mammary stem cell function by steroid hormone signalling. Nature. 2010;465(7299):798–802. Epub 20100411. doi: 10.1038/nature09027. PubMed PMID: 20383121.

10. Brisken C, O’Malley B. Hormone action in the mammary gland. Cold Spring Harb Perspect Biol. 2010;2(12):a003178. Epub 20100825. doi: 10.1101/cshperspect.a003178. PubMed PMID: 20739412; PubMed Central PMCID: PMCPMC2982168.

11. Joshi PA, Waterhouse PD, Kasaian K, Fang H, Gulyaeva O, Sul HS, et al. PDGFRalpha(+) stromal adipocyte progenitors transition into epithelial cells during lobulo-alveologenesis in the murine mammary gland. Nat Commun. 2019;10(1):1760. Epub 20190415. doi: 10.1038/s41467-019-09748-z. PubMed PMID: 30988300; PubMed Central PMCID: PMCPMC6465250.

12. Shehata M, Teschendorff A, Sharp G, Novcic N, Russell IA, Avril S, et al. Phenotypic and functional characterisation of the luminal cell hierarchy of the mammary gland. Breast Cancer Res. 2012;14(5):R134. Epub 20121022. doi: 10.1186/bcr3334. PubMed PMID: 23088371; PubMed Central PMCID: PMCPMC4053112.

13. Kasemeier-Kulesa JC, Morrison JA, Lefcort F, Kulesa PM. TrkB/BDNF signalling patterns the sympathetic nervous system. Nat Commun. 2015;6:8281. Epub 20150925. doi: 10.1038/ncomms9281. PubMed PMID: 26404565; PubMed Central PMCID: PMCPMC4586040.

14. Sar Shalom H, Goldner R, Golan-Vaishenker Y, Yaron A. Balance between BDNF and Semaphorins gates the innervation of the mammary gland. Elife. 2019;8. Epub 20190110. doi: 10.7554/eLife.41162. PubMed PMID: 30628891; PubMed Central PMCID: PMCPMC6328272.

15. Brisken C. Progesterone signalling in breast cancer: a neglected hormone coming into the limelight. Nat Rev Cancer. 2013;13(6):385–96. doi: 10.1038/nrc3518. PubMed PMID: 23702927.

16. Joshi PA, Goodwin PJ, Khokha R. Progesterone Exposure and Breast Cancer Risk: Understanding the Biological Roots. JAMA Oncol. 2015;1(3):283–5. doi: 10.1001/jamaoncol.2015.0512. PubMed PMID: 26181171.

17. Folkerd E, Dowsett M. Sex hormones and breast cancer risk and prognosis. Breast. 2013;22 Suppl 2:S38–43. doi: 10.1016/j.breast.2013.07.007. PubMed PMID: 24074790.

18. Colitti M. Expression of NGF, BDNF and their high-affinity receptors in ovine mammary glands during development and lactation. Histochem Cell Biol. 2015;144(6):559–70. Epub 20150823. doi: 10.1007/s00418-015-1360-0. PubMed PMID: 26298090.

19. Ge Y, Cao Y, Zhang J, Li F, Wang J, Sun M, et al. GOS enhances BDNF-mediated mammary gland development in pubertal mice via the gut-brain axis. NPJ Biofilms Microbiomes. 2024;10(1):130. Epub 20241119. doi: 10.1038/s41522-024-00607-4. PubMed PMID: 39562762; PubMed Central PMCID: PMCPMC11577074.

