## Supplementary Information for "A sex hormone-BDNF-TrkB axis directs sympathetic innervation in the mouse mammary gland"

### Supplementary figure 1, Maity et al.

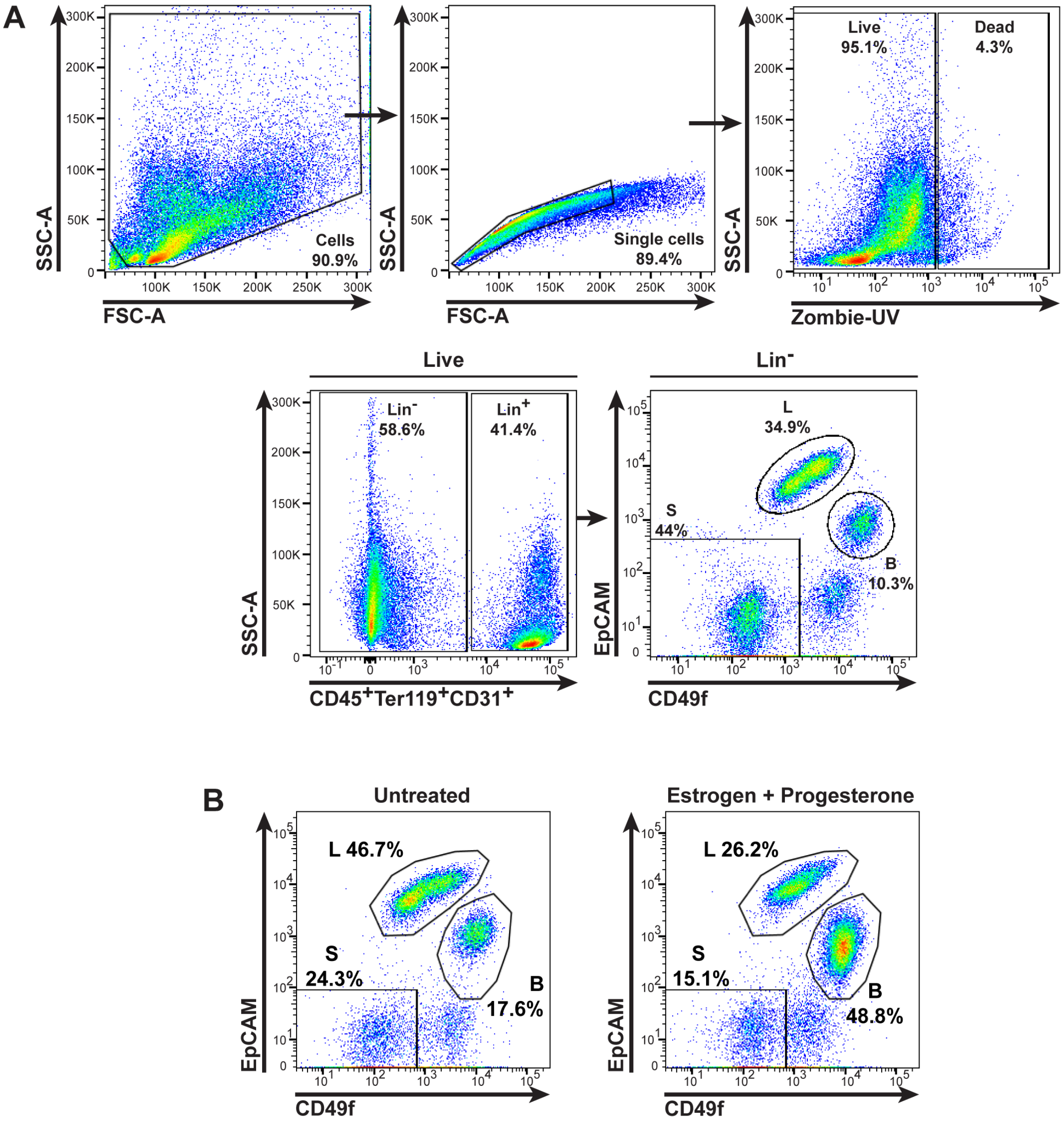


**Supplementary Figure 1: Fluorescence-activated cell sorting of mammary luminal epithelial cells.**

**(A)** Gating strategy to sort mammary luminal epithelial cells. **(B)** Representative plots showing live lineage (lin)-negative cells of untreated vs estrogen + progesterone (EP)-treated samples.

**Supplementary Table**

| Antibodies | Vendor | Catalog number | Working concentration |
|---|---|---|---|
| Anti-Human PGP 9.5 (UCHL1) (184), (affinity purified) (Rabbit IgG) | Cedarlane | CL7756AP-50 | 1:1000 (IF) |
| Anti-Tyrosine Hydroxylase Antibody | Sigma | AB152 | 1:400 (IF) |
| Human/Mouse E-Cadherin Antibody | R&D Systems | AF748 | 1:200 (IF) |
| Invitrogen BDNF Polyclonal antibody | Fisher | PIPA585730 | 1:50 (IF) |
| Invitrogen™ TrkB Polyclonal Antibody | Fisher | PIPA578405 | 1:50 (IF), 1:1000 (WB) |
| Invitrogen™ Phospho-TrkB (Tyr516) Polyclonal Antibody | Fisher | PIPA536695 | 1:1000 (WB) |
| Cy™3 AffiniPure® Donkey Anti-Rabbit IgG (H+L) | Cedarlane | 711-165-152 | 1:300 (IF) |
| Alexa Fluor® 647 AffiniPure® Donkey Anti-Goat IgG (H+L) | Cedarlane | 705-605-003 | 1:300 (IF) |
| Cy™3 AffiniPure® Donkey Anti-Rat IgG (H+L) | Cedarlane | 712-165-150 | 1:300 (IF) |
| Anti-rabbit IgG, HRP-linked Antibody | Cell Signaling Technology | 7074S | 1:3000 (WB) |
| PE-Cy7 anti-CD45 antibody | Fisher | 25-0451-82 | 1:1000 (FACS) |
| PE-Cy7 anti-Ter119 antibody | Fisher | 25-5921-82 | 1:200 (FACS) |
| PE-Cy7 anti-CD31 antibody | Fisher | 25-0311-82 | 1:500 (FACS) |
| ZombieUV viability dye | BioLegend | 423107 | 1:100 (FACS) |
| APC-Cy7 anti-EpCAM antibody | BioLegend | 118218 | 1:200 (FACS) |
| APC anti-CD49f antibody | R&D Systems | FAB13501A | 1:50 (FACS) |
